# Cellular pathways of calcium transport and concentration towards mineral formation in sea urchin larvae

**DOI:** 10.1101/2020.08.17.244137

**Authors:** Keren Kahil, Neta Varsano, Andrea Sorrentino, Eva Pereiro, Peter Rez, Steve Weiner, Lia Addadi

## Abstract

Sea urchin larvae have an endoskeleton consisting of two calcitic spicules. The primary mesenchyme cells (PMCs) are the cells that are responsible for spicule formation. PMCs endocytose sea water from the larval internal body cavity into a network of vacuoles and vesicles, where calcium ions are concentrated until they precipitate in the form of amorphous calcium carbonate (ACC). The mineral is subsequently transferred to the syncytium, where the spicule forms. Using cryo-soft X-ray microscopy (cryo-SXM) we imaged intra-cellular calcium-containing particles in the PMCs and acquired Ca-L_2,3_ X-ray absorption near edge spectra (XANES) of these Ca-particles. Using the pre-peak/main peak (L_2_’/ L_2_) intensity ratio, which reflects the atomic order in the first Ca coordination shell, we determined the state of the calcium ions in each particle. The concentration of Ca in each of the particles was also determined by the integrated area in the main Ca absorption peak. We observed about 700 Ca-particles with order parameters, L_2_’/ L_2_, ranging from solution to hydrated and anhydrous ACC, and with concentrations ranging between 1-15 M. We conclude that in each cell the calcium ions exist in a continuum of states. This implies that most, but not all water, is expelled from the particles. This cellular process of calcium concentration may represent a widespread pathway in mineralizing organisms.

**Significance:** Organisms form mineralized skeletons, many of which are composed of calcium salts. Marine organisms extract calcium ions from sea water. One of the main unresolved issues is how organisms concentrate calcium by more than 3 orders of magnitude, to achieve mineral deposition in their skeleton. Here we determine the calcium state in each of the calcium-containing vesicles inside the spicule-building cells of sea urchin larvae. We show that within one cell there is a wide range of concentrations and states from solution to solid. We hypothesize that calcium concentration increases gradually in each vesicle, starting from sea water levels and until mineral is deposited. This model might well be relevant to other phyla, thus advancing the understanding of biomineralization processes.

## Introduction

Calcium ions play a critical role in many cellular processes. Calcium ions are messengers for a wide range of cellular activities, including fertilization, cell differentiation, secretion, muscle contraction and programmed cell death (1, 2). Therefore, the concentration of Ca^2+^ in the cytosol is highly regulated at around 100-200 nM during resting stages (3–5). Ca^2+^ homeostasis is mediated by Ca-binding proteins and different organelles in the cell, including the endoplasmic reticulum (ER) and the mitochondria that serve as significant Ca^2+^ stores and as signal generators (6–8).

Calcium ions are an important component of many biominerals such as bones, teeth, shells and spines (9). In comparison to Ca^2+^signaling, which requires small amounts of Ca^2+^, biomineralization processes require massive sequestering and transport of ions from the environment and/or from the food, to the site of mineralization. The sequestered ions can reach the mineralization site as solutes, but can also concentrate intracellularly inside vesicles, where they precipitate to form highly disordered mineral phases (10–15). In the latter case, specialized cells take up the ions through ion pumps, ion channels, or by endocytosis of extracellular fluid, and process the calcium ions until export to the final mineralization location (16–19).

In this study, we evaluate the contents of calcium-containing vesicles in primary mesenchyme cells (PMCs), which are involved in the formation of the calcitic skeleton of the sea urchin larvae. In this way, we obtain insights into how calcium ions, extracted from the environment, are concentrated and stored for spicule formation.

*Paracentrotus lividus* sea urchin embryos form an endoskeleton consisting of two calcitic spicules, within 72 hours after fertilization (20, 21). The source of the calcium ions is the surrounding sea water, whereas the carbonate ions are thought to originate from both sea water and metabolic processes in the embryo (22, 23). Sea water enters the embryonic body cavity (blastocoel) through the permeable ectoderm cell layer of the embryo (24). Endocytosis of sea water and blastocoel fluid into PMCs, as well as endothelial and epithelial cells (25, 26) was tracked by labeling sea water with calcein, a fluorescent calcium-binding and membrane impermeable dye (17, 26–28). The endocytosed fluid in the PMCs was observed to form a network of vacuoles and vesicles (26). Beniash et al. observed electron-dense granules of sizes 0.5-1.5 μm in PMCs, which are composed of amorphous calcium carbonate (ACC) (10). Intracellular vesicles of similar size were observed Vidavsky et al., using cryo-SEM and air-SEM, containing calcium carbonate deposits composed of nano-particles, 20-30 nm in size (25). Ca-deposits within the same range of sizes were also observed in the rough ER of PMCs (29).

The intracellularly produced ACC is subsequently exported to the growing spicule, where it partially transforms into calcite through secondary nucleation (30–33). The location and distribution of ACC and calcite in the spicule were studied by using extended X-ray absorption fine structure (EXAFS), X-ray absorption near edge spectroscopy (XANES) and photoelectron emission microscopy (PEEM) (30, 34, 35). Three distinct mineral phases were identified in the spicule: hydrated ACC (ACC*H_2_O), anhydrous ACC and crystalline calcite (34). According to theoretical simulations of Rez and Blackwell (36), the two different amorphous phases arise from different levels of ordering of the oxygen coordination polyhedron around calcium. The coordination polyhedron becomes more ordered as the transformation to the crystalline phase progresses. The phase information contained in the XANES spectra is exploited here to characterize the mineral phases in the intra-cellular vesicles.

Cryo-soft X-ray transmission microscopy (cryo-SXM) is an attractive technique for tomography and spectromicroscopy of biological samples in the hydrated state (37–39). Imaging is performed in the “water window” interval of X-ray energies, namely between the carbon (C) K-edge (284 eV) and the oxygen (O) K-edge (543 eV). As a result, in the “water window” C is highly absorbing, whereas O and thus H_2_O, is almost transparent. Subsequently, carbon-rich moieties such as lipid bodies, proteins and membranes appear dark in transmission, whereas the water rich cytosol appears lighter (40). The Ca L_2,3_-absorption edge between ~346 and 356 eV also resides in the “water window” (41). Therefore, imaging across this absorption edge enables the characterization of Ca-rich moieties in whole, hydrated cells. Each pixel of the same field of view, imaged as the energy is varied across the Ca L_2,3_-edge, can be assigned an individual Ca L_2,3_-edge XANES spectrum. This technique was applied to the calcifying coccolithophorid alga (42, 43). In this study, we use cryo-SXM and XANES to locate and characterize both the phases and the concentrations of Ca-rich bodies in sea urchin larval cells.

## Results

The intracellular calcium contents of sea urchin larval PMCs, as well as non-PMC cells, were studied using cryo-SXM. Cells were obtained from disaggregated embryos 36 hours post fertilization (hpf), when the spicule building activity is maximal (20, 44). A fluorescent dye that specifically labels PMCs was added to the cell suspension (45, 46). The labelled cell suspension in sea water was seeded onto gold finder-TEM-grids and the grids were vitrified by plunge freezing. The grids were imaged using cryo-fluorescence and bright field microscopy in order to locate the labeled PMCs (*Materials and Methods* and *SI Appendix*, Fig. S1).

### Cryo-SXM imaging of cells

Tomograms of PMCs and non-PMCs of sea urchin embryos were acquired first by performing tilt series at 352.6 eV, namely the Ca L_2_-edge energy (Movie S1). Due to the attenuation of the X-rays, information can be acquired from 15 μm thick samples, allowing imaging of complete cells disaggregated from embryos. After reconstruction, different cellular compartments with high contrast relative to the cytoplasm are visible (Movies S2 and S3). These include the cell membrane (M), the nucleus (N) and different vesicles and vacuoles (V), which may contain both C and Ca. Both PMCs and non-PMCs contain large organelles with medium contrast, but PMCs show in addition a large number of very high contrast bodies that are 100-500 nm in size (Movies S2-S3, Dark Bodies – DB, Table S1). We then collected 2D projection images of 9 PMCs and 12 non-PMCs at the Ca L_2_-edge energy, 352.6 eV, where Ca absorbs the most (Fig. 1*A* and *E*), and below the edge at 338.3 eV (Fig. 1*B* and *F*). Since the absorption from the other elements is almost the same at these two energies, subtracting the images taken below the edge from the images taken at the Ca edge results in 2D “Ca-maps”, which show the locations of concentrated Ca (Fig. 1*C* and *G*, white areas).

**Fig. 1.**
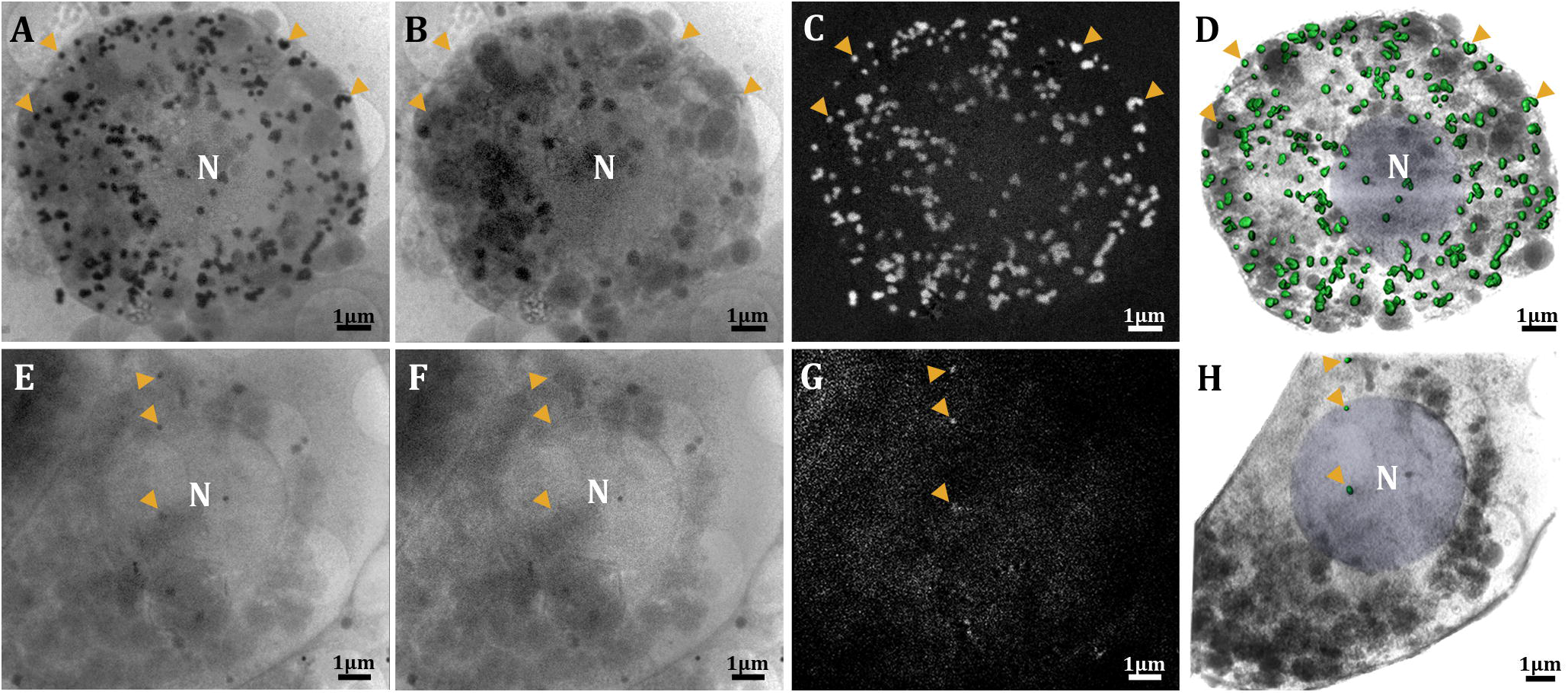
Identification of Ca-particles using cryo-SXM in a PMC (*A-D*) and a non-PMC (*E-H*). (*A* and *E*) 2D projection transmission images of the PMC and the non-PMC (respectively) taken at the Ca L_2_-edge (352.6 eV), (*B* and *F)* 2D projection transmission images of *A* and *E*, respectively, taken below the Ca L_2,3_-edge (338.3 eV), where Ca is almost transparent. Orange arrowheads point to selected particles that are not visible in *B* and *F* but are visible in *A* and *E*, respectively. These are Ca-particles. (C) and (G) “Ca-maps” obtained by subtraction of *B* from *A* and *F* from *E*. The grey scale is converted to absorbance, so that white areas are locations where Ca is concentrated. (*D* and *H)* Superimposition of the 2D Ca-particle maps on the 3D segmented and reconstructed tomograms of both cells. Green – Ca-particles, blue – nucleus (N), grey – cell membranes, cytoplasm, vesicles and vacuoles that do not contain Ca.

The Ca-maps show that PMCs contain many Ca-bearing bodies, whereas non-PMCs contain few Ca-bearing bodies (Fig. 1 *A-C*, *E-G*). This was the case for all PMCs and non-PMCs examined (*SI Appendix*, Fig. S2 and Table S1). We refer to these Ca-bearing bodies as Ca-particles. The 2D Ca-maps were then superimposed on the 3D reconstructed data from the tomograms of the same PMC and non-PMC cells. The segmented Ca-particles could, in this way, be differentiated from other cellular organelles (Fig. 1 *D, H* and Movie *S4*, Ca-particles in green).

The Ca-particles can be found in several distribution patterns: isolated (Fig. 2*A*, inset), several particles clustered in one larger body (Fig. 2*B*, inset), or one particle contained in a large vesicle (Fig. 2*C*, inset). The lack of a detectable membrane around the isolated Ca-particles in Fig. 2*A*, and the poorly defined membrane visible in Fig. 2*B*, may be due to the high absorbance of calcium at the Ca L-edge masking the membrane in contact with the Ca-particles, or to the lower carbon contrast at those energies compared to that obtained when imaging at 520 eV (*SI Appendix*, Fig. S3). These images are similar to observations of vesicles in PMCs made with cryo-SEM (Fig. 2 *D, E*). In cryo-SEM, membrane-bound vesicles of 0.5-1 μm in size are observed to contain many aggregated particles of ≈20 nm size (Fig. 2*D*). Alternatively, denser particles with sizes 200-500 nm are observed within vesicles (Fig. 2*E*).

**Fig. 2.**
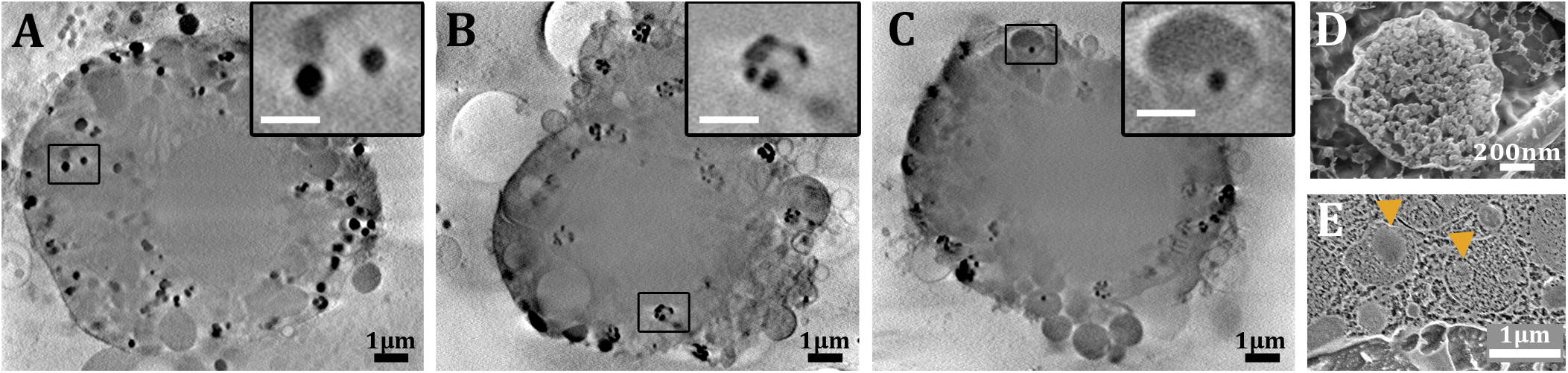
Ca-particle distribution patterns. (*A-C*) Slices through the reconstructed volumes of 3 PMCs. Insets: high magnifications of the boxed areas (*A*) Ca-particles not enveloped by a visible membrane. (*B*) Clusters of Ca-particles within a larger body. (*C*) Ca-particle contained in a larger vesicle. (*D, E*) Cryo-SEM micrographs of vesicles from sea urchin embryos. The vesicles are inside PMCs. Scale bar in the insets=0.5 μm. (*D*) Vesicle from high pressure frozen and freeze fractured embryo. The vesicle contains granular, 20 nm particles. (*E*) Vesicles from a high pressure frozen and cryo-planed embryo. The vesicles contain denser particles (arrowheads) surrounded by medium with a texture similar to the cytoplasm.

### Cryo-SXM Ca edge spectroscopy: phase characterization

The presence of abundant Ca-particles inside the PMCs raises questions regarding the nature of the phases of Ca in these particles. Phase information can be obtained from the XANES spectra of each Ca-particle (Fig. 3A). The ratio of the L_2_ pre-peak (L_2_’) to the L_2_ peak (L_2_’/L_2_) is a good indication of the short-range order of the oxygen atoms around the calcium ions (30, 35, 36). In general, the higher the ratio of L_2_’/L_2_, the more ordered the phase is. The spectra of calcite, anhydrous ACC and hydrated ACC (ACC*H_2_O) taken from (35), are used here as references for the known calcium carbonate phases in the sea urchin larval spicule (Fig 3A, black, green and purple curves). The values of L_2_’/L_2_, measured by XPEEM, are: ACC*H_2_O L_2_’/L_2_= 0.26, anhydrous ACC L_2_’/L_2_= 0.4 and calcite L_2_’/L_2_= 0.45 (Fig. 3A). The sea water XANES spectrum, measured by cryo-SXM, has no L_2_’ peak (Fig. 3A, blue curve) in accordance with theory (36).

**Fig. 3.**
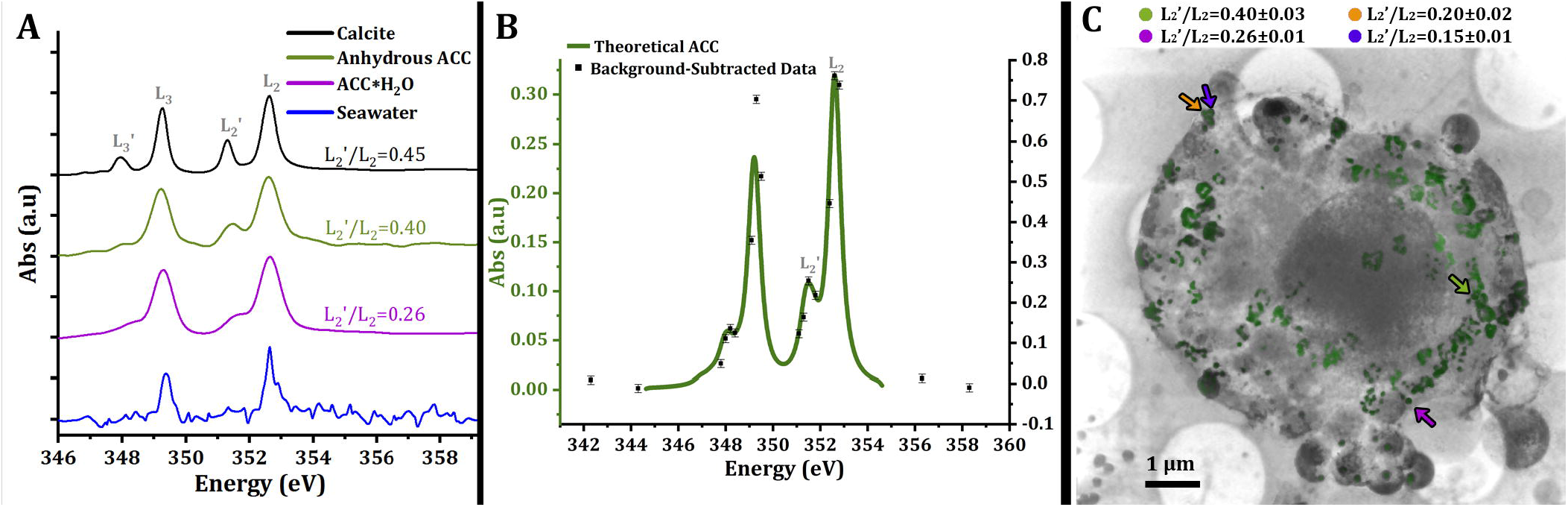
Ca L_2,3_ X-ray Near-Edge Spectroscopy (XANES). (*A*) XANES of four standard samples: Calcite in black, anhydrous ACC in green, hydrated ACC (ACC*H_2_O) in purple (35) and sea water in blue. (*B*) XANES spectrum obtained for one particle, from the measurement of the 18 data points, after background subtraction. Green spectrum = ACC calculated spectrum from (36). The procedure is explained in detail in *SI Appendix, SI Materials and Methods*, XANES). (C) Superimposition of the 2D Ca-particle map on the 3D reconstructed and segmented PMC cell tomogram. The data is overlaid on a slice through the reconstructed volume. Green – Ca-particles, grey – cell membranes. Four particles (colored arrows) with their L_2_’/L_2_ values are shown.

To retrieve the individual XANES spectra of Ca-particles in a cell, ideally images of the whole cell are recorded as the energy is varied in small steps across the Ca L_2,3_-edge, from 338.3 – 358.3 eV. In this way, a full XANES spectrum is obtained from each pixel in the image (pixel size=13nm). In order to acquire such spectra with good signal-to-noise ratios and spectral resolution, an exposure time of several seconds is needed at each energy step. Empirically we observed that after about 20 energy steps, visible damage could be seen. We consequently selected 18 energy points, placed along the Ca L_2,3_-edge at key energy values, in such a way as to still enable characterization of the L_2_’ pre-peak and L_2_ peaks (Fig. 3*B*), without causing damage *(Materials and methods* and *SI Appendix*, Fig. S4). The average absorbance at each energy value of each Ca-particle was obtained by averaging the signal at that value of all the pixels in the same Ca-particle. The Ca-particle area was determined according to the Ca-maps. To illustrate the analytical approach, Fig. 3*B* shows the data obtained for one particle, from the measurement of the 18 data points, after background subtraction. Peak-intensity ratios of L_2_’/L_2_ were calculated using the background-subtracted absorbance of L_2_’ (absorbance of L_2_’ at 351.5eV) and L_2_ (absorbance of L_2_ at 352.6eV). The parameter dominating the uncertainty of the data is the background fitting error, which in turn is dominated by the counting error *(SI appendix, Materials and methods*). Fig. 3C identifies 4 Ca-particles inside a PMC. Two of the Ca-particles have L_2_’/L_2_ values similar to anhydrous ACC and to ACC*H_2_O (Fig. 3C green and purple points, respectively), whereas the magenta and blue particles are more disordered than any known amorphous phase (Fig. 3C).

We are aware that the absorbance signal for the L_2_ peak may reach saturation, resulting in an over-estimation of L_2_’/L_2_. In addition, as the XANES data points are averaged across a particle, the L_2_’/L_2_ values may result from a linear combination of several states within one particle. To address these issues we plotted the L_2_-peak intensity value vs the L_2_’-peak intensity value of the raw data of each pixel from 40 randomly selected particles. If the Ca state across the particle is homogeneous and the L_2_ peak signal does not reach saturation, we expect a linear relation between L_2_’ and L_2_ peak intensities. Three categories of particles were observed *(SI Appendix*, Fig. S5): (1) linear relation between the peak intensities without saturation, (2) a scattered relation and (3) a linear relation that reaches saturation (Fig. S5 *A, B* and *C*, respectively). We observed these 3 categories for both small particles (hence few pixels) and large particles (hence many pixels). The relative scatter in the linear relation is within the uncertainty of the L_2_/L_2_’ value. About 20% of the examined particles contained saturated pixels. However, the number of saturated pixels is relatively small in each particle (<5%) and is therefore within the uncertainty of the L_2_’/L_2_ value. Particles in which the scatter in L_2_’/L_2_ is greater than the uncertainty are considered to be heterogeneous, i.e. having different states.

We determined the L_2_’/L_2_ values for around 700 Ca-particles in 5 different PMCs, and observed that we cannot classify the Ca-particles into distinct groups with defined disorder. The results of analyzing more than 400 Ca-particles from two representative cells (Fig. 4*Ai*, *Aii*) are shown in Fig. 4*B*. There is a continuum of disordered states starting from very low values of L_2_’/L_2_, similar to sea water, and ranging to values similar to anhydrous ACC (Fig. 4*B*). Very few Ca-particles (5 particles out of >700) show an L_2_’/L_2_ value similar to calcite, within experimental error. Around 50% of the Ca-particles have L_2_’/L_2_ values between anhydrous ACC and ACC*H_2_O and the other 50% have L_2_’/L_2_ values lower than ACC*H_2_O, i.e. with even lower short-range order.

**Fig. 4.**
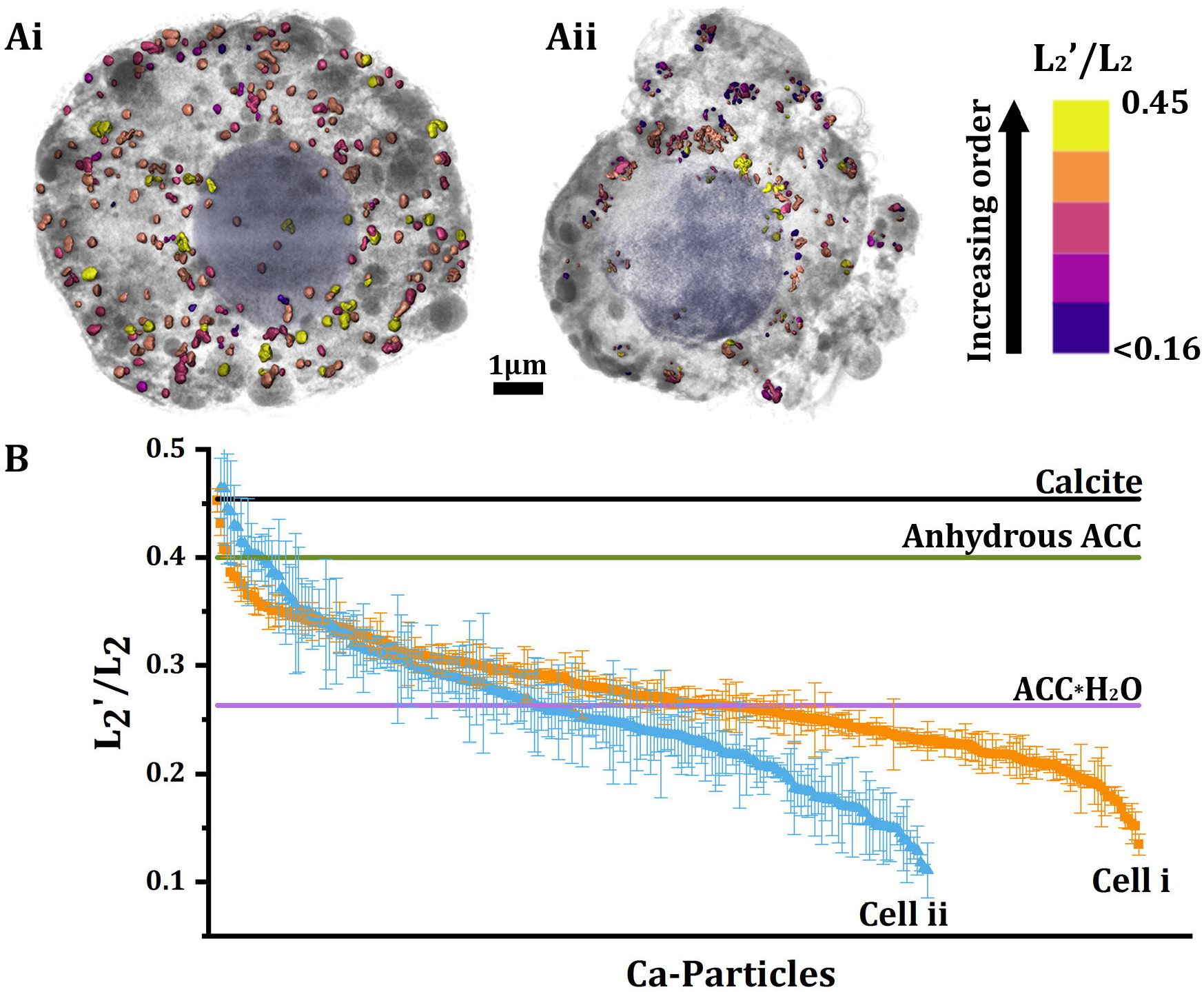
Ca-particle L_2_’/L_2_ value distributions. (A*i*, A*ii*) two segmented reconstructed volumes of PMCs tomograms. Ca-particles are marked in colors according to their L_2_’/L_2_ value, as indicated in the color scale on the right side. Blue – nucleus, grey – cytoplasm, membranes and C-rich vesicles and vacuoles. The average diameter of the Ca-particle in the two cells are: for cell i =230±50 nm, and for cell ii = 180±70 nm. (*B*) L_2_’/L_2_ values for all the Ca-particles in the 2 cells shown in A (cell i in orange and cell ii in blue). The values are sorted for convenience in descending order to show the general trend. There is no correlation between the L_2_’/L_2_ values and the location of a particle within the cell as shown in *A*.

The observed distribution of the L_2_’/L_2_ values in Fig. 3 might raise legitimate concerns that the data represent a random distribution of errors around the ratio of ACC*H_2_O. To address these concerns, we calculated the curves corresponding to random distribution of L_2_’/L_2_ values with similar errors for the two cells (*SI Appendix*, Fig S6). The simulated curves were based on average errors of 5% and 10% for Cell i, which has a calculated average error of 4%, and for Cell ii 10% and 20%. Cell ii has a calculated average error of 10%. The simulated curves are much narrower than the observed distributions for both cells, confirming that the measured data represent a true distribution of states.

The Ca-particles were segmented and digitally colored according to their L_2_’/L_2_ values (Fig. 4*A*i, ii). We observed that Ca-particles with different levels of disorder can be in close proximity or possibly even connected (Movie S5). In fact, there is no discernible Ca-particle distribution pattern inside the volume of the cell (Movie S5), although we note that the spatial distribution may have changed during disaggregation.

### Calcium concentrations in the Ca-particles

The X-ray absorption of each Ca-particle at the Ca absorption edge is proportional to the Ca concentration in the particle. Thus, the calcium concentration can in principle be extracted from the spectroscopic data, provided the X-ray absorption cross section of Ca is known, and assuming linearity between X-ray absorbance and Ca concentration (for more details see *Materials and Methods*). The Ca absorption cross section was theoretically estimated to be σ= 0.0625 Å^2^ per atom (*SI Appendix, Materials and Methods*). For each Ca-particle, we estimated the integrated absorption by fitting a Gaussian to the L_2_-peak data points. The effective thickness was taken as 0.67 of the particle diameter, to account for the particles being spherical (based on the 3D reconstructed data of the cells). The concentration assessment was performed only for particles located within the 2 μm of the cell closest to the beam entry surface (Fig 5A, colored green). This procedure was adopted because, for a particle located in depth inside the cell, the attenuation of the beam before encountering the particle may substantially affect the absorption of the particle itself. Note that the beam intensity absorbed by the cell before or after the beam hits the particle is the same as in the background, and is subtracted as background. Small particles with diameters <100nm were not taken into consideration, to avoid large relative errors in the evaluation of their sizes. The error was calculated based on the error in the peak-fitting coefficients (*Material and Methods*).

**Fig. 5.**
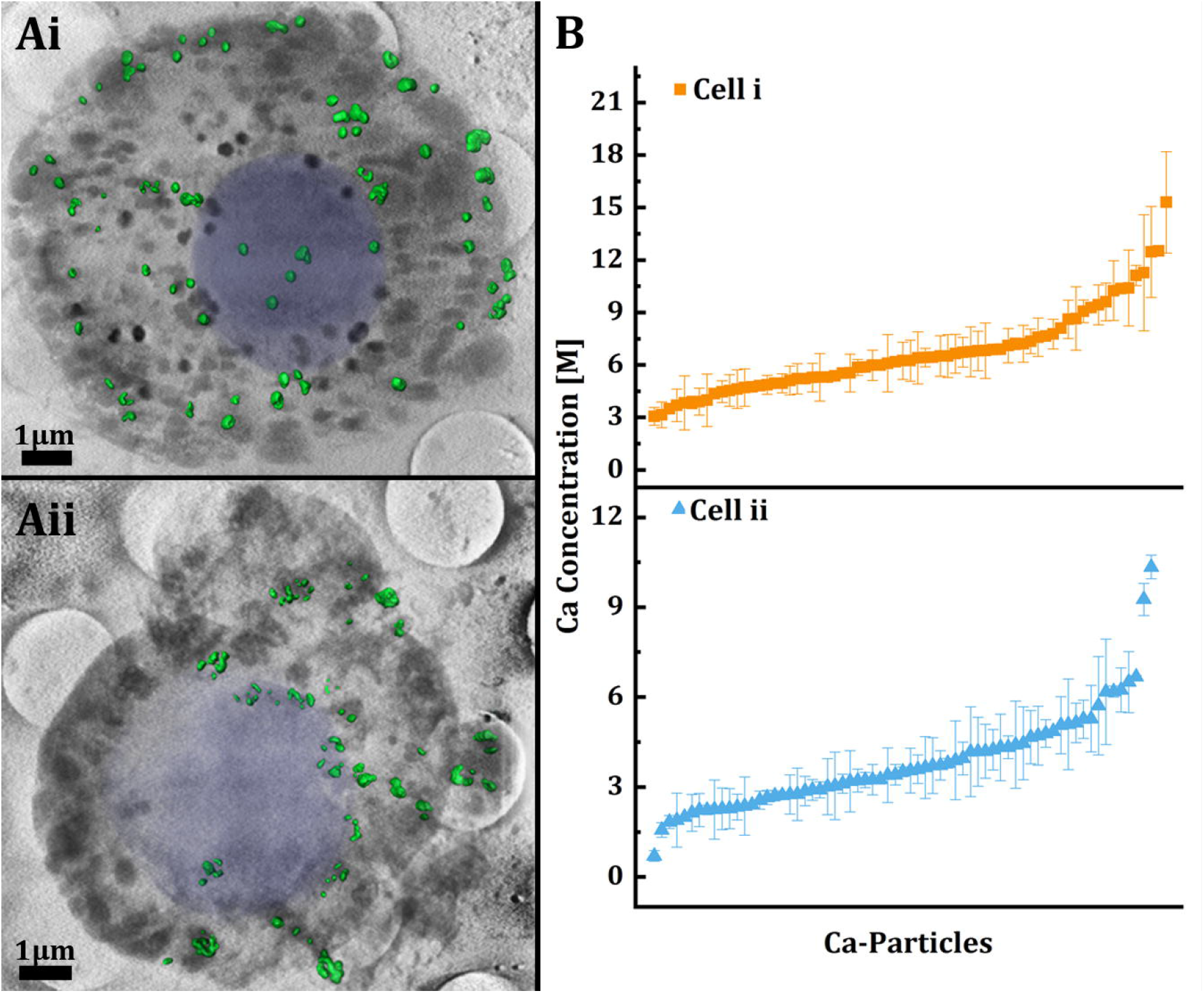
Ca concentration estimation. (*Ai and Aii*) Superimposition of the cell segmentation on the 3D reconstruction of the PMCs presented in Fig. 4. In green are the Ca-particles within 2 μm of the cell closest to the beam entry surface, which were used for the concentration calculations (see text). (*B*) Ca concentrations of the particles in the cells shown in *Ai* and *Aii*. The values are sorted in ascending order for convenience. Please note that the concentrations may be slightly underestimated, because the area of the L_2_’ peak was not taken into account in the integrated absorption of the L_2_ peak. The extent of the underestimation falls, in any case, within the limits of the error.

The Ca concentration values for 140 Ca-particles in two cells are shown in Fig. 5*B*. There is no discernible clustering of the values, which range continuously from ~1M up to ~15M. Note that the estimated minimum concentration detection limit, for particles within the range of sizes in Table S1, is ~1M (for more details see *SI Appendix, Materials and Methods*). All the values are much higher than sea water (10mM Ca), but still lower than calcite, which is 27M, and lower than ACC*H_2_O, which is 19M.

As mentioned above, saturation of the signal can result in decreased L_2_-peak intensity, resulting in an under-estimation of the concentration. We note, however, that the uncertainty of the concentration values is larger than the effect of the few saturated pixels (less than 5%) on the concentration. A weak correlation with Pearson correlation coefficients of 0.03 for cell i and 0.08 for cell ii, was observed between the concentration values and the L_2_’/L_2_ values, i.e. the degree of order (*SI Appendix*, Fig. S7).

## Discussion

We observe that the PMCs of sea urchin embryos contain tens to hundreds of dense Ca-bearing particles, 100-500 nm in size, whereas non-PMCs contain very few, and in some cases no particles. The short-range order of the oxygen atoms around the calcium ion show that approximately half of the Ca-particles have L_2_’/L_2_ values smaller than those of hydrated ACC, indicating an even less ordered state. The remaining half of the Ca-particles have L_2_’/L_2_ values between those of hydrous and anhydrous ACC, in a continuous range of values between the two known ACC phases. The estimated concentrations of calcium in these Ca-particles range from 1 M to 15 M. Based on these observations, we discuss processes occurring between sea water endocytosis from the blastocoel fluid into PMCs (26), and the final deposition of calcium carbonate mineral into the spicules (10, 30).

The source of ions for spicule deposition is sea water (22, 23). In sea water, calcium is found in concentrations of about 10 mM, and carbonate, which is ~6% of the dissolved inorganic carbon at the sea water pH of 8.2, is around 0.3 mM (47, 48). Therefore, sea water is already supersaturated by >3 orders of magnitude relative to calcite (K_sp_= 3.3×10^-9^ (49)) and one order of magnitude relative to ACC (K_sp_=4×10^-7^ (50)). Calcium carbonate should therefore form, but its formation is presumably inhibited by other ions and molecules in the sea water (51, 52).

Sea water first enters the larval body cavity, the blastocoel. The calcium concentrations in the blastocoel are presumably similar to the concentration in sea water, as inferred from the permeability and lack of pH regulation of the ectoderm layer (24). Larvae grown in sea water containing calcein, take up the calcein first into the blastocoel and then into vesicles within PMCs, epithelial cells and endothelial cells (25). As calcein does not pass through membranes, this uptake process must involve endocytosis (17). We cannot exclude the possibility that additional calcium and/or carbonate ions are introduced into the vesicles through selective ion pumps, but we note that this is not essential.

Sea water enters the PMCs by endocytosis in large vacuoles and vacuolar networks, which are then transformed into vesicles (26). Our cryo-SXM observations show that different PMCs have different numbers of Ca-containing particles and that the levels of short-range order, as well as Ca concentrations, vary continuously within a PMC. We therefore infer that the sea water in these vesicles undergoes continuous and gradual changes in composition, including removal of different ions and of water. We note that the two PMC cells analyzed in detail have different ranges in the number and in the state of the calcium particles. This may reflect different stages of PMC involvement in spicule formation, depending on the cell location in the larva and on the developmental stage.

The calcium concentrations increase in the vesicles from that of sea water (10 mM) up to ~15 M. It is plausible that under these conditions of increasing calcium and presumably carbonate concentration, calcium carbonate will form from dissolved calcium ions in the calcium rich particles. Even though the solubility of different calcium carbonate species is different (53–55), and is certainly influenced by the local composition of ions and organic components in the vesicle, within this range of concentrations we expect that the initial liquid phase will transform into a solid.

Although we cannot follow the time-evolution of the Ca-particles, it is conceivable that our observation of a continuum of short-range order levels around the calcium ion reflects the process of ACC deposition. The fact that the degree of order changes in a continuum, indicates that there are no abrupt phase transitions of whole Ca-particles. Furthermore, Ca L-edge XANES is not specifically sensitive to the counter ion and we cannot determine if and at what stage the counter ion is carbonate. We note, however, that Naftel et al (56) recorded Ca L-edge XANES spectra for a series of salts with different counter-ions, including carbonate, phosphate, chloride and several organic molecules. These spectra show substantial shifts in the positions of the L_2_ and L_2_’ peaks, and more importantly substantial differences in the ΔeV= eV_L2_-eV_L2’_. As the L_2_ and L_2_’ values measured here are the same as those measured for calcium carbonates, Naftel’s data support the notion that the particles measured in the PMCs, albeit at Ca concentrations > 1M, have carbonate counter ions. It is reasonable to assume that Ca-particles with very low levels of order correspond to calcium ions in solution, whereas Ca-particles with L_2_’/L_2_ values similar or above hydrated ACC are solid (36, 57).

We have observed vesicles filled with 20 nm particles by cryo-SEM, and vesicles containing larger and denser particles, which significantly do not fill the whole vesicle volume. These different mineral-containing vesicles may indicate two successive stages of mineral condensation (58). The maximum calcium concentration measured by cryo-SXM (15 M), however, is lower than the concentration of hydrated ACC (19 M), even taking into account a slight underestimation in the values reported in Fig. 5. This gap in concentrations may be due to the conceivable co-existence of two types of water in close proximity to the CaCO_3_ – liquid water and structural water. The 20 nm particles are presumably formed of hydrated ACC, i.e. contain structural water, and are separated by liquid water. Even the larger and denser particles, although they appear to be full of mineral, are probably composed of smaller units, such that there is still plenty of space for liquid water.

It is plausible that ACC aggregates, with associated residual ions and/or organic molecules, leave the PMCs and reach the spicule with a large excess of water within them. Only upon attachment to the spicule, followed by fusion and crystallization, would the water be removed. The structural water content may be further regulated by Mg^2+^ ions, which have a high propensity to be hydrated. The excess Mg ions would then be expelled together with the water of hydration, upon crystallization.

Many open questions remain. The calcium concentration process described here does not span the whole concentration range of calcium from sea water to ACC. It would be interesting to follow the initial stages of calcium concentration from sea water (0.01 M) to the minimum concentration detected by cryo-SXM in this study (~1 M).

Another interesting question concerns the involvement of the other sea water ions, primarily Na, Cl and Mg. How and when are they removed in the process of CaCO_3_ precipitation and mineralization? The carbonate pathway remains an unsolved and highly intriguing issue. When does the calcium we observe change from calcium in solution to CaCO_3_?

In sea water, the carbonate concentration is much lower than the calcium concentration. If the pH in the vesicles increases to 9, as was observed for the unicellular foraminifera (17), the carbonate concentration would increase from 0.3mM to 1.1mM. Alternatively, the CO_3_^2-^ concentration will increase if the CO_2_ and/or HCO_3_^-^ concentrations increase in the vesicle (59). In addition, it is suggested that the conversion of CO_2_ and HCO_3_^-^ to CO_3_^-2^ in the sea urchin larva is mediated by carbonic anhydrase (60, 61). Interestingly, no carbonic anhydrase activity was found in foraminifera, which undergo a similar process of carbonate concentration (62).

Calcium plays an important role in all cells, functioning as first and second messenger for a variety of cell processes (1). Yet, calcium can also lead to cell apoptosis or necrosis if its cytosolic concentration, which is 100–200 nM under normal conditions, is not properly controlled (63). Therefore, it is not surprising that the reported calcium stores, including sea water inclusions, inside cells, are compartmentalized inside membrane-delimited spaces (17, 25, 26, 64). Similarly, although we do not observe membranes around all the Ca-particles, it follows from the above that the Ca-particles must be membrane bound and the whole process of mineral formation occurs in a membrane-delimited space.

Sea urchin embryos of *Paracentrotus lividus* at 36hpf (prism stage), the age studied here, are in the prime of spicule building. Thus, it is not surprising that the PMCs, the cells responsible for mineral deposition, are full of Ca-rich particles. However, less than 10 Ca-particles per cell are present in non-PMCs, compared to hundreds of Ca-particles in PMCs. Previous studies of sea urchin embryos observed calcein-labeled vesicles in some ectoderm cells in similar amounts as in the PMCs (25). While tracking the fate of endocytosed sea water into vesicles, Vidavsky et al observed that incorporated dextran and calcein were co-localized in almost 100% of the vesicles in epithelial cells (26). In PMCs however, more than 40% of the cells were labeled only with calcein. It is conceivable that the vesicles labeled exclusively with calcein in PMCs are the ones that accumulate mineral and therefore correspond to the vesicles we observe using cryo-SXM. The vesicles labeled with calcein in the epithelial cells may reflect a calcium concentration <1M, the minimum concentration detected by cryo-SXM in this study.

Little is known to date about intra-cellular ion transport and concentration on the pathway towards mineral deposition in most organisms. The pioneering studies of Bentov, Erez et al first showed sea water endocytosis and processing toward mineralization in foraminifera (17). Intra-cellular vesicles containing amorphous mineral were observed in bone-forming cells in zebrafish (65), in mice (66) and in chicken embryos (67). Furthermore, the involvement of intra-vesicular amorphous calcium phosphate and amorphous calcium carbonate as precursor phases of many biogenic minerals was reported (68–70). It is noteworthy that incubation with calcein results in the staining of the skeletal minerals not only in foraminifera and in echinoderms, but also in cnidarians, arthropods, molluscs, brachiopods and chordates (71). Following Erez, we conclude that in these organisms there is direct communication and transport of calcium in sea water to the site of mineralization. To date, only in coccolithophorids and dinoflagellates, was it demonstrated that incubation with calcein does not result in labeling of the newly deposited mineral, indicating a different ion transport pathway (42, 71, 72). Here we show that in the PMCs of echinoderms intracellular ion concentration occurs within individual vesicles and reaches a point where mineral forms and can then be exocytosed into the spicule-forming syncytium. This concentration process may also be relevant to mineralizing cells in other phyla.

## Materials and Methods

### Sea urchin larval culture

Mature cultured adult *Paracentrotus lividus* sea urchins were supplied by the Israel Oceanographic and Limnological Research Institute, Eilat. Spawning, fertilization, and embryo development were carried out as described (73).

### Sample preparation for cryo-SXM

Sea urchin embryos at 36 hours post fertilization were chemically and mechanically disaggregated following the method of Giudice and Mutolo (74). Wheat Germ Agglutinin-rhodamine (RL-1022, Vector Laboratories, Inc.) was added to the cell suspension at a concentration of 5 μg/mL, and left for 5 min, in order to specifically label the PMCs (45). The suspension was then centrifuged at 900 rcf for 5 min. The cell pellet was kept and diluted with 5 mL of filtered fresh sea water (from the Red Sea in Eilat).

The cell suspension in sea water was seeded onto R 2/2 (SiO_2_) on Au G200F1 finder grids (Quantifoil Micro Tools GmbH, Germany) and incubated with filtered fresh sea water at 18 °C overnight. For plunging, the grids were lifted and held by tweezers inside the plunge freezer chamber at 90% relative humidity and 22 °C (Leica EM-GP plunger, Leica Microsystems). 1 μL drop of X5 concentrated gold nanoparticle (150 nm) suspension (746649, Sigma Aldrich), was added to each grid to provide fiducial markers for tomography. The grids were blotted for 3 s and plunge frozen into liquid ethane.

Even though the cells are collected from embryos 36 hours post fertilization, they are cultivated overnight before cryo-fixation. During this time, the PMCs are lacking the signaling factors that are usually provided by cells of the ectoderm, and their absence most probably interrupts the spicule formation activities (75, 76). If signaling factors are present, the cells may be more developed than 36 hours.

Calcite sample preparation: pure calcite crystals (Iceland spar) were finely ground into powder using a mortar and pestle. 100 μL of DDW were added to 2 mg calcite powder and mixture was stirred. Immediately after mixing, 4 μL of the suspension was placed on QUANTIFOIL^®^ R 2/2 Cu 200 grid, blotted for 3s and plunge frozen.

Sea water liposome preparation: 50 μL of egg chicken L-α-phosphatidylcholine (95%) in chloroform, 25 mg/mL (131601C, MERCK) were placed in 1 mL centrifuge tube and left to evaporate under a N_2_ stream. 1 mL of fresh sea water was added to the tube and vortexed for 20s to create multi-lamellar liposomes. 4 μL of the liposome suspension was placed on QUANTIFOIL^®^ R 2/2 Cu 200 grid, blotted for 3 s and plunge frozen.

All samples were kept in liquid nitrogen until loaded at the synchrotron beam line microscope.

### Cryo-fluorescence and bright field microscopy imaging

Following plunge freezing, cell-containing grids were loaded onto a cryo-correlative microscopy stage (CMS196M, LINKAM SCIENTIFIC INSTRUMENTS, UK) and imaged both in bright field and in fluorescence mode using a 515-545nm LED for excitation and a TRITC filter for collection. Images were analyzed using ImageJ (US National Institutes of Health, Bethesda, Maryland, USA). Spherical cells with sizes >6μm and emitting a fluorescent signal were identified as PMCs (45, 46) (*SI Appendix*, Fig. S1).

### Soft X-ray cryo-tomography and spectromicroscopy

X-ray imaging was performed at the MISTRAL beamline (ALBA Synchrotron, Barcelona, Spain) (77). Initially, a tilt series at 352.6 eV X-ray energy was collected to allow 3D volume reconstruction of cells and their internal structures. At this energy, corresponding to the Ca L_2_-edge peak maximum, Ca-rich bodies are highly absorbing. The tilt series consisted of 121–131 images taken at 1° degree intervals. Exposure time was 4 s to optimize signal-to-noise level while minimizing radiation damage. No radiation damage was observed using this exposure time at the achieved spatial resolution. The data sets were acquired using a 40 nm zone plate objective lens which give a half pitch resolution of 31 nm (78). The effective pixel size in the projection images was 13nm. The projection images of the tilt series were normalized using the flat-field (average of 10 images with no sample, collected at 2s exposure time), to take into account the intensity distribution delivered to the sample by the capillary condenser lens. The alignment of the tomographic projections was performed in Bsoft (79) using the gold nano-particles, or dark features inside the cells, as fiducial markers. The aligned projection tilt series were reconstructed in TomoJ using the ART algorithm with 15 iterations and 0.1 relaxation coefficient (80). Visualization and segmentation of the final volumes were carried out using the Avizo 9.5 3D software (Thermo Fisher Scientific, USA). The Ca-particles were individually identified using the Ca-maps (mentioned below) and automatically segmented.

Energy scan series around the Ca L_2,3_-edge of standard samples was acquired by imaging the same field of view at varying X-ray energies. The acquisition procedure was as follows: in the range 338.3-342.3 eV with 0.25 eV steps, 342.3-354.3eV with 0.1eV steps and 354.3-358.3 eV at 0.5 eV steps. Each image was taken with 3s exposure time. The energy scan of calcite was taken as a reference for the Ca L_2,3_ energies. All the energy values where shifted in the text to have the L_2_ peak maxima at 352.6 eV following Gong et al. (35).

To obtain better signal-to-noise ratios and reduce beam damage to the cells, 18 energies were selected: 342.3 eV and 344.3 eV in the pre-edge region; 347.8 eV, 348 eV and 348.2 eV on the L_3_ pre-peak (L_3_’); 348.4 eV between L_3_’ and L_3_; 349.1 eV, 349.3 eV and 349.5 eV on the L_3_ peak; 351.1 eV, 351.3 eV and 351.5 eV on the L_2_’ peak, 351.8 eV between L_2_’ and L_2_; 352.4 eV, 352.6 eV and 352.8 eV on the L_2_ peak; and 356.3eV and 358.3eV in the post edge region. During the energy scans the objective zone-plate lens and the charge-coupled device detector positions were automatically adjusted to maintain focus and constant magnification. Five images at 2s exposure/image were taken at each energy and averaged. To estimate the counting error at a specific energy (352.6 and 351.5eV), we calculated the standard deviation (STD) of the counts from the five images taken at that energy for each pixel. We then calculated the averaged STD for the pixels in each Ca-particle. The averaged counting error of the Ca-particles is ~10% of the counts at both energies. The averaged image was normalized using the flat-field (one image with no sample collected at 2s exposure time). The procedure used to evaluate the counting error of the flat-field image is described in SI (Fig S10).

### XANES analysis

Each normalized stack of transmission images acquired in the energy scan was transformed from transmission to absorbance by taking the (–) natural logarithm of each image using ImageJ (81). Calcium 2D localization in the samples was carried out by subtracting an image of the cell taken at 338.3 eV (before the Ca L-edge) from the image taken at 352.6 eV (Ca L_2_-edge). The “Ca-maps” for all cells were examined, and areas of Ca signal were marked and saved. XANES spectra of the Ca-particles were extracted for every Ca-particle by plotting the averaged absorbance of all the pixels in the marked particle area, as a function of the energy at which the image was taken. The spectra were then subjected to background subtraction. Detailed explanations and error estimation are reported in *SI appendix, Materials and methods*.

### Ca concentration estimation

Concentration calculations were carried out assuming linearity between absorbance, *A*, and Ca concentration using *A*=*μt*, where *A* is the integrated intensity over the L_2_ peak, *μ* is the linear absorption coefficient and t is the thickness. To make sure that the calculated concentration is not affected by absorption of the cell bulk, only Ca-particles found in the 2 μm of the cells closest to the beam entry surface were selected using the reconstructed 3D data. To account for the Ca-particles being spherical, as shown by the 3D reconstructed data, their effective thickness, i. e. the length the beam passed in the particle, was taken as 0.67 of their diameter (see detailed explanation in *SI appendix, Materials and methods*).

The L_2_-peak data points were fitted with a Gaussian and integrated using MATLAB (The MathWorks Inc., USA). The error in the concentration values was estimated as the half range of the 95% confidence interval of the fitting coefficients calculated by MATLAB, and assuming the errors in the coefficients are dependent (see detailed explanation in *SI appendix, Materials and methods*).

### Cryo-SEM freeze fracture and cryo-planing

Sample preparation for cryo-SEM: Embryos ~36 h post fertilization, were immersed in 0.2 mL filtered fresh sea water solution containing 10 wt% dextran (31389, Sigma Aldrich) as a cryo-protectant. 2 μL of the embryo suspension was sandwiched between two metal discs (3-mm diameter, 0.1-mm cavities) and cryoimmobilized in a high-pressure freezing device (HPM10, Bal-Tec AG, Liechtenstein). For cryo-planing one of the metal discs was immersed in hexadecane (H6703, Sigma Aldrich) and removed after high-pressure freezing. The frozen samples were kept in liquid nitrogen.

Freeze fracture samples: The samples were transferred by using a vacuum cryotransfer device (VCT 100; Leica Microsystems, Vienna, Austria) to a freeze-fracture device (BAF 60; Leica Microsystems). Samples were freeze-fractured at −120 °C, etched for 10 min at −105 °C, and coated with 2.5 nm platinum/carbon by double-axis rotary shadowing. Samples were observed at −120 °C in an Ultra 55 SEM (Zeiss, Germany) (25).

Cryo-planed samples: The disc containing the frozen sample was transferred to a cryomicrotome (UC6, Leica Microsystems, Vienna, Austria) and sectioned at −150°C in a nitrogen atmosphere as described by Mor Khalifa et al (82). Samples were observed at −120 °C in an Ultra 55 SEM (Zeiss, Germany).

## Acknowledgments

We thank David Ben-Ezra and Muki Shpigel from the Israel Oceanographic and Limnological Research Center for supplying mature sea urchins and Assaf Gal for inspiration and fruitful discussions. Cryo-SXM experiments were performed at the MISTRAL beamline at ALBA Synchrotron with the collaboration of ALBA staff. This research was supported by the Minerva foundation with funding from the Federal German Ministry for Education and Research. K.K. is the recipient of the Levzion fellowship from the Israeli Council for Higher Education. L.A. is the incumbent of the Dorothy and Patrick Gorman Professorial Chair of Biological Ultrastructure at the Weizmann Institute of Science.

