## Supplementary Information for "Cellular pathways of calcium transport and concentration towards mineral formation in sea urchin larvae"

**This PDF file includes:**

Figures S1 to S6

Legends for Movies S1 to S5

SI Materials and Methods containing figures S7- S10

SI References

**Other supplementary materials for this manuscript include the following:**

Movies S1 to S5

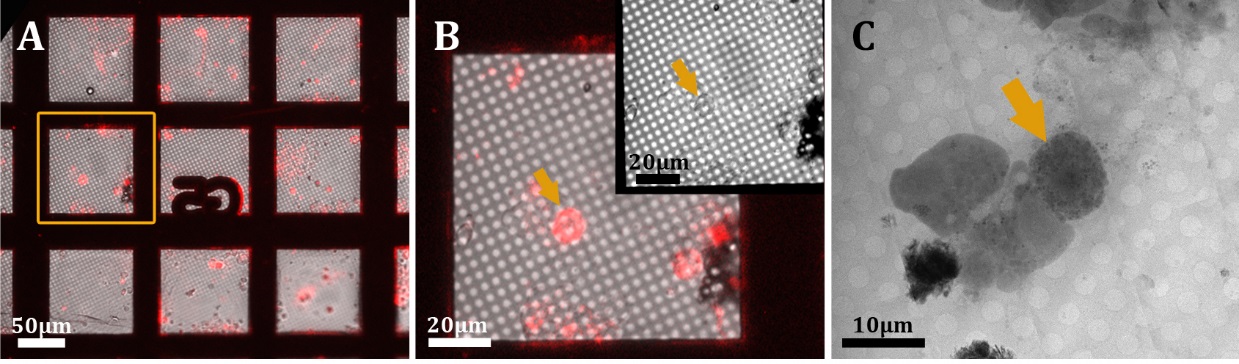

Fig. S1. Identification of a PMC using cryo-fluorescent light microscopy. (*A*) An overview of part of a plunged grid. The image is a superimposition of the bright field channel with the fluorescence channel (Red). Excitation - 515-545 nm, TRITC filter. (*B*) Higher magnification of the boxed square in *A* showing a labeled cell (arrow). Inset – bright field channel of the same field of view. (*C*) The same cell from *B* imaged using cryo-SXM at 352.6 eV.

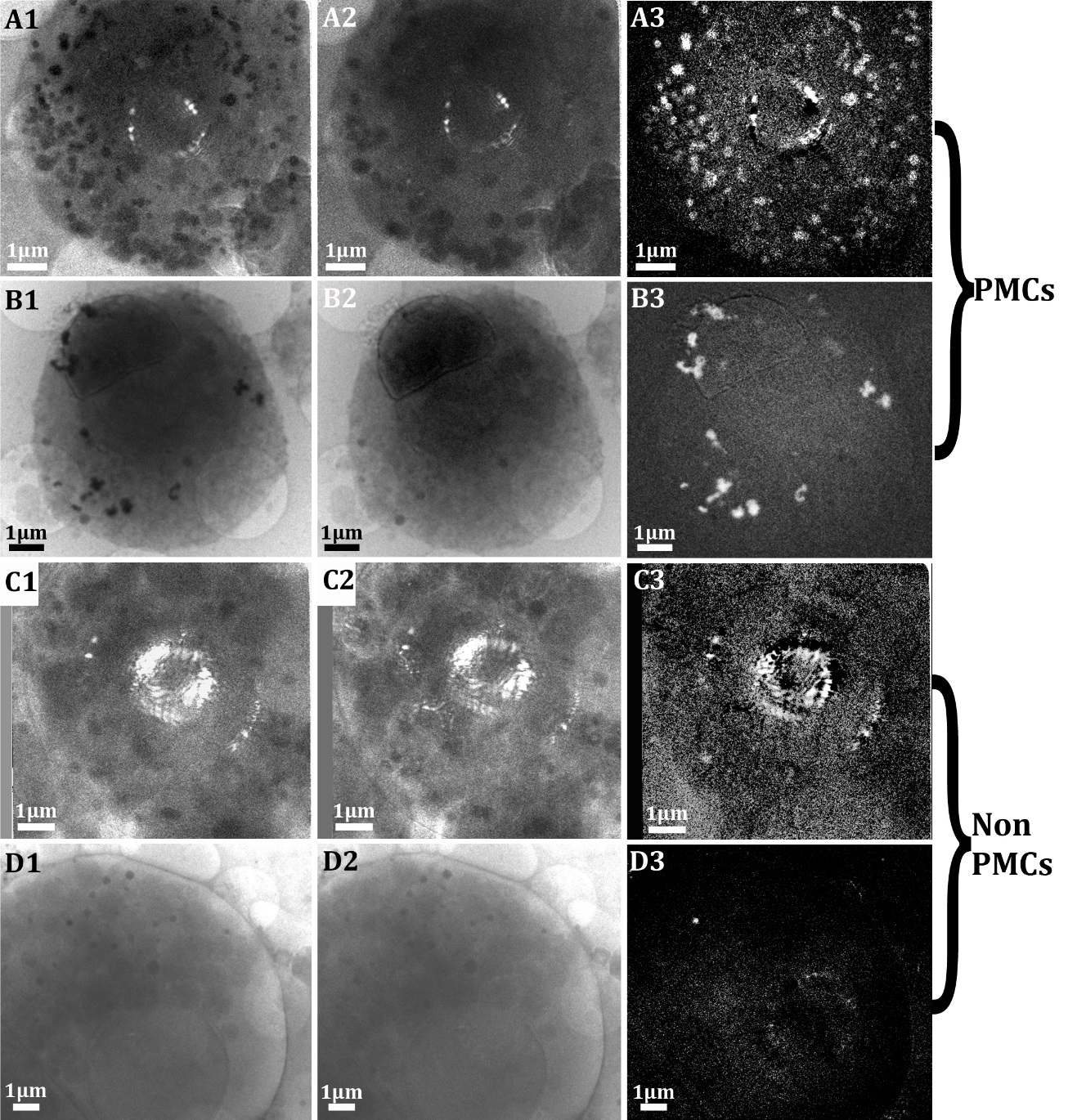

Fig. S2. Identification of Ca-particles using cryo-SXM. (A1, B1) and (C1, D1): 2D projection images of a PMC and a non-PMC (respectively) taken at the Ca L_2_-edge. (A2, B2) and (C2, D2): 2D projection images of a PMC and a non-PMC (respectively) taken below the Ca L_2,3_-edge (344eV), where Ca is almost transparent. (A3, B3) and (C3, D3): “Ca-maps” obtained by subtraction of X2 from X1 (X=A, B, C, D). The grey scale is converted to absorbance, so that white areas are locations where Ca is concentrated. The bright spots that form a circle in the middle of (A), (C) and (D) are direct beam scattering artefacts on the beam stop located before the condenser lens which are visible when the cells are very absorbing (related to cell thickness). The data in (A) and (C) were acquired using a 25 nm zone plate, pixel size 15.8 nm. The data in (B) and (D) were acquired using 40 nm zone plate, pixel size=13 nm.

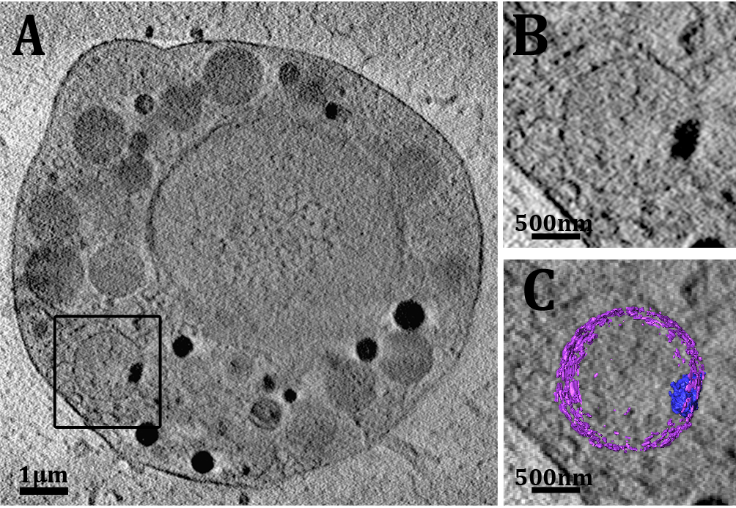

Fig. S3. Cryo-SXT performed at 520 eV. (A) A slice through reconstructed data of a cell disaggregated from a sea urchin larva. The data were acquired at the energy of 520eV, such that the dark particle in the membrane-delineated vesicle (box) may contain both C and Ca. (B and C) Higher magnification of the boxed vesicle in A. (C) Superimposition of B with segmentation of the whole vesicle – 1µm in depth. Purple - vesicle membrane, blue – dark precipitate.

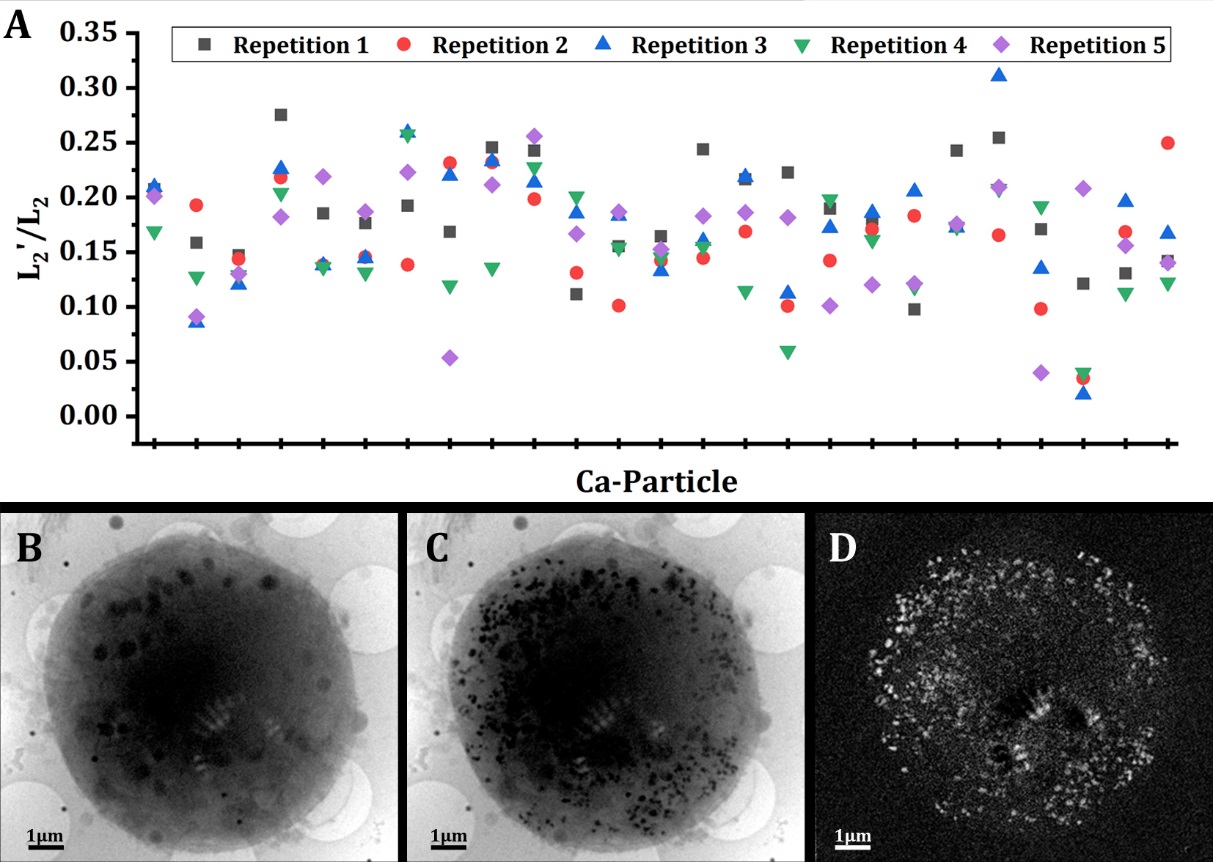

Fig. S4. Radiation damage evolution with time. To exclude radiation damage caused to the Ca-mineral, the phase of the Ca-particles was examined with time. In the measurement series, each absorbance measurement was taken 5 times at each energy, and the average of the 5 measurements was recorded. To evaluate possible radiation damage, one image at 2 s exposure time per energy was acquired for all 18 energy values, and the scan was repeated 5 times. (A) L2’/L2 values for 25 Ca-particles from the cell for each scan repetition (repetition 1-5). There is no increase or decrease in the L2’/L2 from the first repetition to the last in the majority (>90%) of the particles, indicating the absence of radiation damage. (B and C) A PMC imaged below (340 eV) and on the Ca-L_2_ edge respectively. (D) “Ca-map” of the same cell. The map was obtained by subtracting the image in (B) from the image in (C). White dots are areas of concentrated Ca-mineral. The bright spots that form a circle in the middle of (B-D) are direct beam scattering artefacts on the beam stop located before the condenser lens.

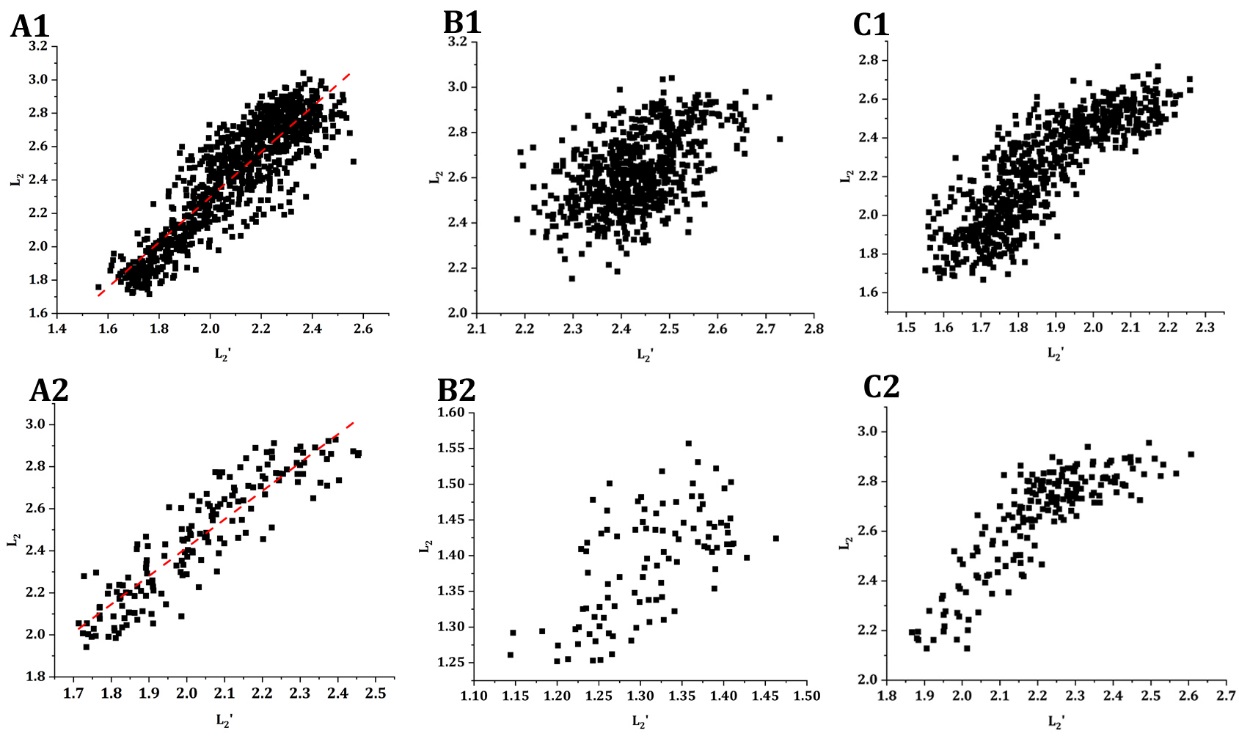

Fig. S5. Pixel-by-pixel plots of the L_2_ peak intensity vs the L_2_’ peak intensity values of the raw data of each pixel from 6 individual particles out of 40 randomly selected particles that were analyzed. (A1 and A2) Large and small particles (respectively) showing a linear correlation between L_2_ intensity and L_2_’ intensity (red dashed line shows the general linear-trend line). No saturation is observed. (B1 and B2) Large and small particles (respectively) showing no correlation, namely a scattered relation, between the peak intensities. (C1 and C2) Large and small particles (respectively) showing a linear correlation between the peak intensities. Both plots show a region where the L_2_’ increases while the L_2_ increases in a non-linear manner – these latter data points are saturated pixels.

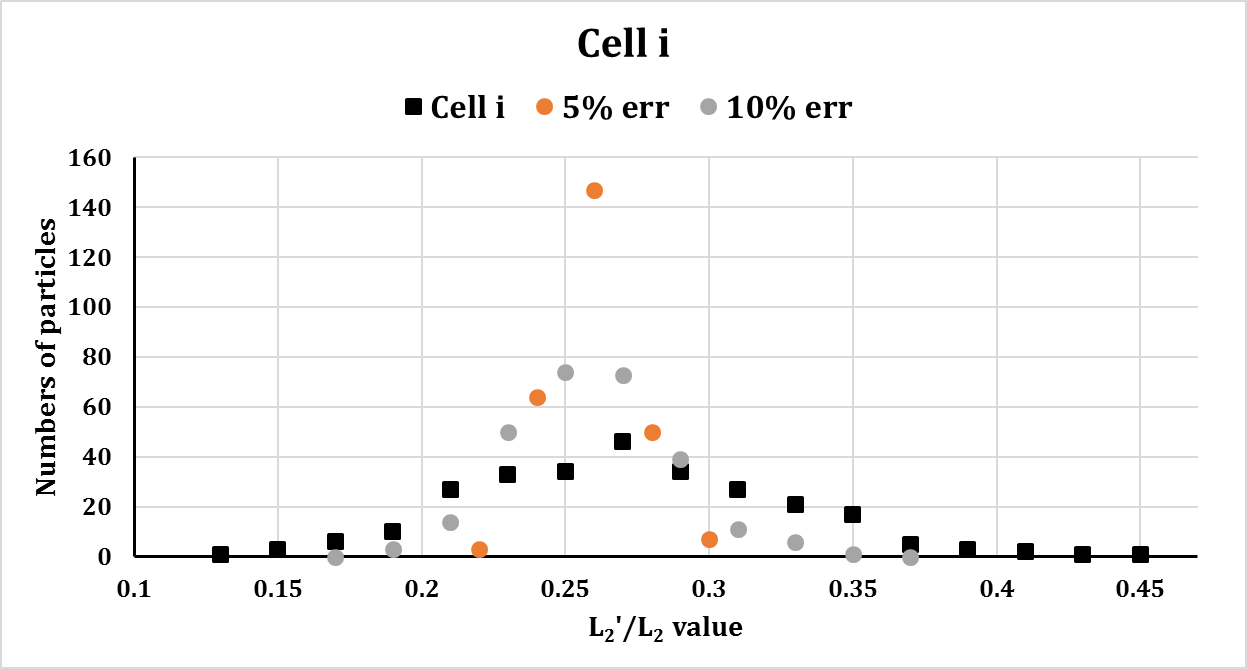

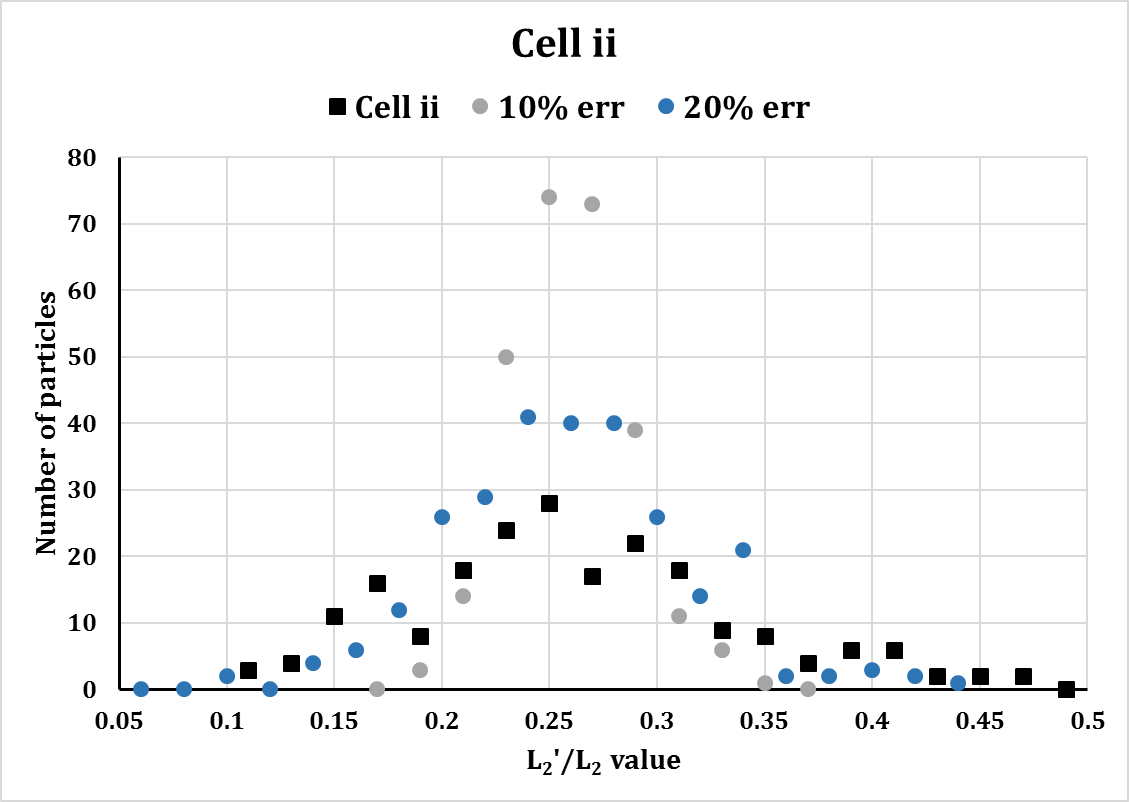

Fig. S6 – Histograms of measured and randomly generated L_2_’/L_2_ values. The random values were generated with normal distribution around L_2_’/L_2_=0.26 (hydrated ACC) and different standard deviations. The full width at half maximum (FWHM) of the randomly generated distributions are: for simulated error 5% - FWHM≈0.07, for simulated error 10% - FWHM≈0.15, for simulated error 20% - FWHM≈0.28. The FWHM of the measured values (presented in fig. 4): for Cell i, which has an averaged error of 4%, FWHM≈0.29 and for Cell ii, which has an averaged error of 10%, FWHM≈0.35. The curves showing the measured data are clearly wider than the simulated curves.

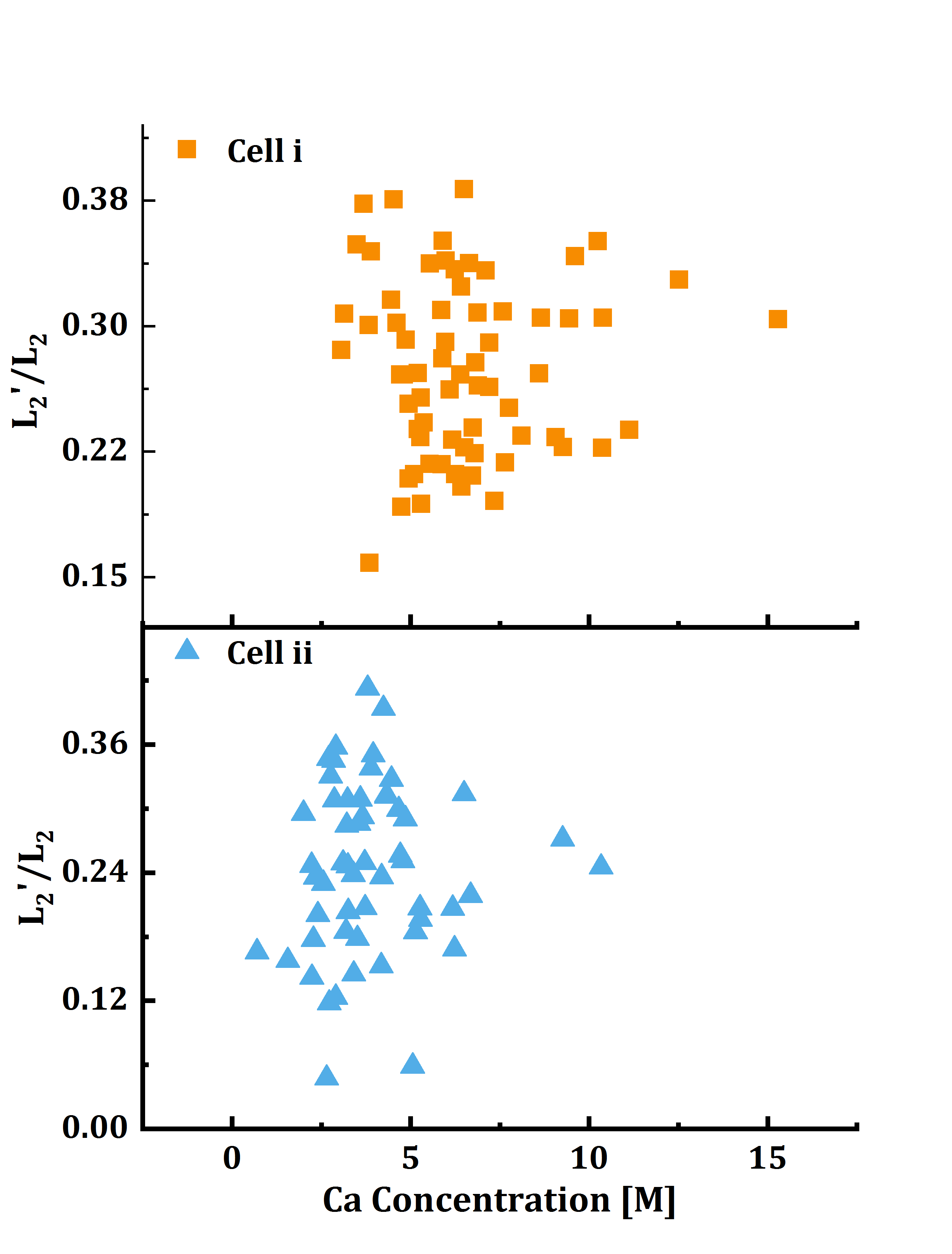

Fig. S7. Plots of Ca concentration vs. order for the cells presented in Figs. 3 and 4 in the main text. Correlation analysis shows a very weak correlation of 0.03 for cell i and 0.08 for cell ii.

Movie S1 (separate file). A tilt series of a PMC (presented in Fig. 1 *A-D*) collected at 352.6 eV X-ray energy using cryo-SXM. The tilt series consists of 131 images taken at 1° intervals. Exposure time=4 s. The data set was acquired using a 40nm zone plate objective lens. The projection images of the tilt series were normalized using the flat-field and aligned using some of the dark features inside the cell as fiducials. Pixel size = 13nm.

Movie S2 (separate file). Cryo-SXT 3D reconstruction of the PMC presented in video S1. The reconstruction was performed using ART algorithm (1) with 15 iterations and 0.1 relaxation coefficient. The video shows the different cellular compartments visible with a spatial resolution of ~50 nm. Pause 1: different cellular compartments marked. M- cell membrane, N – nucleus, V – vesicles and vacuoles and DB - high contrast bodies mentioned in the text. Pixel size =13nm.

Movie S3 (separate file). Cryo-SXT 3D reconstruction of the non-PMC presented in Fig. 1 *E-H*. The reconstruction was performed using ART algorithm (1) with 15 iterations and 0.1 relaxation coefficient. The video shows the different cellular compartments visible with a spatial resolution of ~50 nm. Pause 1: different cellular compartments marked: M- cell membrane, N – nucleus and V – vesicles and vacuoles. Pixel size =13nm.

Movie S4 (separate file). Cryo-SXT 3D segmentation of the reconstruction from Supplementary video 2. Pause 1 – The whole segmented cell. Green – Ca particles, blue – nucleus and grey - cytoplasm, vesicles and vacuoles that do not contain Ca, and cell membranes.

**Movie S5 (separate file).** Ca-particles segmentation according to the level of order. The green particles shown at the beginning of the video are all the Ca particles also shown in Supplementary video 4. The particles then re-appear in groups according to their L_2_’/L_2_ values in descending order.

SI Materials and Methods

Ca-particle determination

The Ca-particle locations were determined using the Ca-maps obtained by subtracting the cell image taken at the L_2_’ energy (351.5eV), from the image taken at the L_2_ energy (352.6eV).

The diameter of each particle was determined by locating the particle in both the Ca-map and the reconstructed 3D data from the tomogram of the same cell (taken at the L_2_ energy). The particles were delimited in the XY plane with a circle and the region of interest was measured using Fiji.

**Table S1. Quantification of Ca-particles and Ca-particle diameter in PMC and non-PMC cells**

Although there seem to be a difference in particle size between PMC and non-PMC cells, the number of particles in the non-PMC cells is too small to allow us to reach any definitive conclusion

| Cell Type | Number of particles | Average Diameter |
| --- | --- | --- |
| PMC | 273 | 232±54nm |
| PMC | 18 | 165±32nm |
| PMC | 303 | 180±70nm |
| PMC | 256 | 163±53nm |
| Non-PMC | 5 | 200±61nm |
| Non-PMC | 1 | 390 |
| Non-PMC | 7 | 390±110nm |
| Non-PMC | 3 | 250±10nm |

XANES

The extraction of the XANES spectra was performed on the normalized and averaged stack. This stack contains one image per energy. The data points in the spectra represent the averaged absorbance of the region-of-interest at each energy (Fig. S8, black squares).

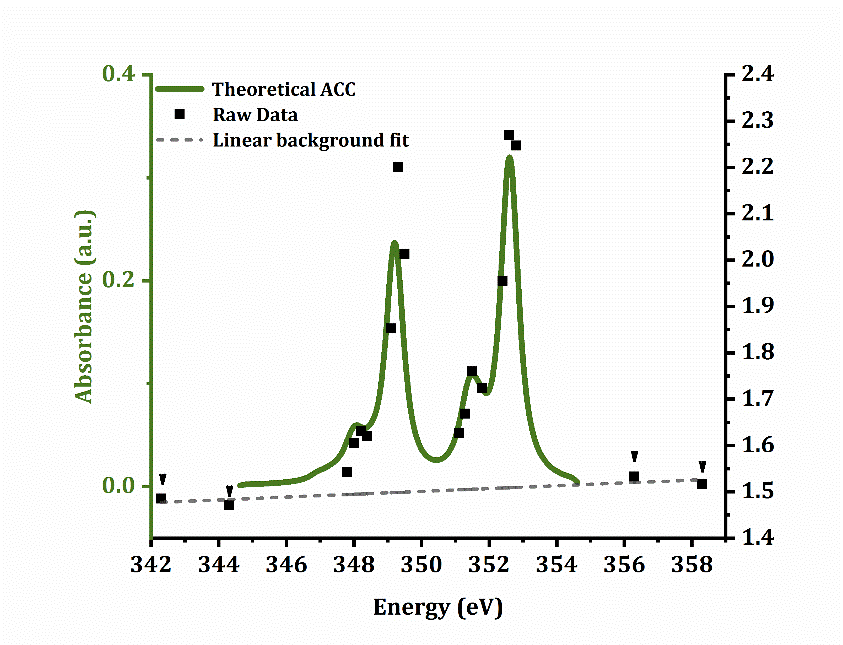

Fig S8. An example for a XANES data set. The averaged and normalized raw data from one particle is shown in black squares. Four data points, two in the pre-edge region and two at the post-edge region, were selected for a linear background fit (marked with arrowheads). The linear background is shown by the dashed grey line. The green full spectrum is theoretically calculated ACC (2).

Background subtraction: A linear trend line was fitted to the following four energy points: 342.3 eV, 344.3 eV 356.3eV and 358.3eV. (Fig. S8, 4 points marked with arrowheads. Dashed grey line = linear fit). The deviation of the 4 points from the fitted line is referred to as ΔI. The linear background was subtracted from the raw data points (Fig. S8), yielding absorbance values for L_2_’ and L_2_. Only spectra with L_2_’ peak intensity/ ΔI > 3 were used throughout the calculations in the text.

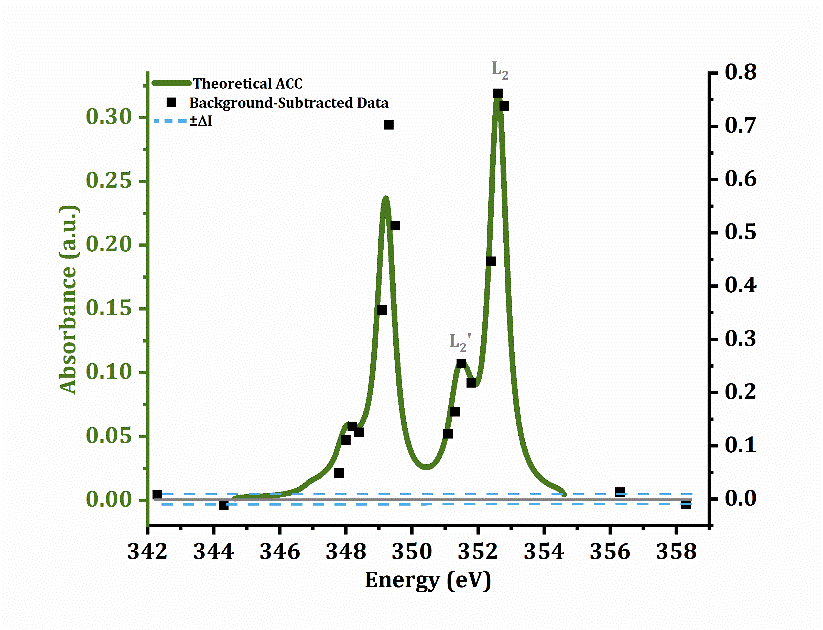

Fig S9. Background subtracted data. The same set of data from Fig. S8 after background subtraction. Blue dashed line show ±ΔI relative to the background, which is now=0. Green spectra – theoretical ACC (2).

L_2_’/L_2_ ratio calculation and error estimation Peak-intensity ratios were calculated using the background-subtracted intensities of L_2_’ (I_L2’_ at 351.5eV) and L_2_ (I_L2_ at 352.6eV).

The error in the I_L2’_/I_L2_ value was estimated considering that the main source of error is in the background estimation:

$$\frac{\left( I_{L2'}\pm\Delta I \right)}{\left( I_{L2}\pm\Delta I \right)}=\frac{I_{L2'}\left( 1\pm\frac{\Delta I}{I_{L2'}} \right)}{I_{L2}\left( 1\pm\frac{\Delta I}{I_{L2}} \right)}=\frac{I_{L2'}}{I_{L2}}\left( 1\pm\frac{\Delta I}{I_{L2'}}\mp\frac{\Delta I}{I_{L2}} \right)$$

The fractional error in the ratio is $f=\Delta I\left| \frac{1}{I_{L2'}}-\frac{1}{I_{L2}} \right|$

The reported error is $E=f\times\frac{I_{L2'}}{I_{L2}}$

We tested the validity of the approach above relative to Gaussian fitting of all four peaks using a Gaussian fitting code. We compared the uncertainties obtained for a few spectra, characterized either by small peaks on a large background or by large peaks on a small background. The estimated errors obtained applying the two procedures were practically the same. This can be explained by considering that for both cases, namely large peaks on a small background (presumably particles near the surface) or small peaks on a high background, the background fitting error dominates. For a particle near the beam entry surface, the background is small, but so is the counting error due to the higher fluence. For particles at greater depths, the background is larger. In either case the background fitting error dominates the counting error.

To estimate the counting error of the flat-field image (I_0_), the square root of the intensity in each pixel was calculated. The fractional counting error of the flat-field image is <0.6% and thus is negligible (Fig S10).

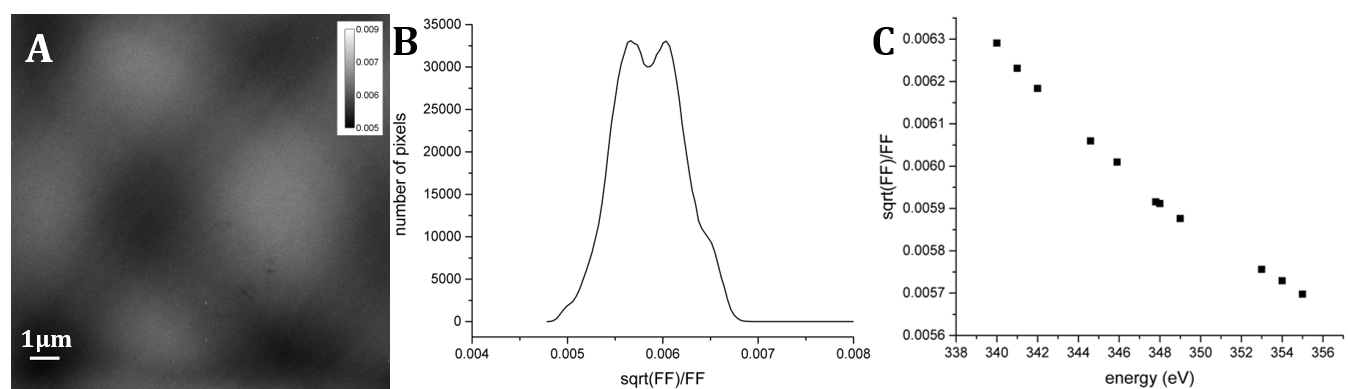

Fig S10. Counting error of the flat-field (FF). (A) calculated standard deviation/FF intensity at 349 eV. (B) Histogram of (A) showing the distribution of relative error in the FF. (C) The averaged relative error in the FF of different energies at the Ca L-edge showing that it remains below 0.6% for all energies.

Concentration Calculation

**Detection limit:** The estimated detection limit is ~1M, based on the following considerations: during calibration measurements, Ca in sea water (10mM = 4x10^-4^ g/cm3) was detected from a 120 nm x 120 nm x 5 µm volume. The particles measured here have a diameter within a range of sizes 150-250nm. If sea water represents the limit of detection in a homogeneous volume, the limit for particles of these sizes should be 600-1000mM (0.6-1M).

To account for the Ca-particles being spherical, as shown by the 3D reconstructed data, their effective thickness, i. e. the length the beam passing through the particle, was taken as 0.67 of their diameter. This effective diameter is the result of projecting a sphere in 2D:

A volume of a sphere is 4/3 πR3. The projected area is πR2, consequently the average thickness of a particle is 4/3 R=2/3 D =0.67 D.

**Linear absorption coefficients:** The linear absorption coefficients, *σ*, are the product of the concentration in Ca atoms per unit volume, *n,* and the absorption cross section, *σ*.

$\mu=n\sigma$	1

The theoretical absorption cross section, σ, was calculated using the theory originally outlined by Cooper (1962)(3), where transitions from all 2p to 3d states were considered, instead of transitions to the continuum.

$\sigma=\frac{480\pi\alpha a_{0}^{2}}{9}\Delta ER^{2}$	2

where *α* is the fine structure constant, *a_0_* is the Bohr Radius, *ΔE* is the energy of the L_2_ peak in Rydbergs and *R* is the matrix element

$R=\int_{0}^{\infty} P_{2p}\left( r \right)rP_{3d}\left( r \right)dr$	3

where $P_{nl}\left( r \right)$ are solutions of the radial Schrodinger equation

$\left( \frac{d^{2}}{dr^{2}}+V\left( r \right)+E_{nl}-\frac{l\left( l+1 \right)}{r^{2}} \right)P_{nl}\left( r \right)=0$	4

The matrix elements were calculated from Dirac-Salter wave functions (4). We estimated, from the observed L_3_:L_2_ ratio, that half the cross section given by equation 2 can be attributed to the L_2_ transition.

**Calculations of the area below L_2_**: The L_2_-peak data points at energies: 352.4eV, 352.6eV, 352.8eV, 356.3eVand 358.3eV (black arrows, Fig. S11), were fitted using MATLAB (The MathWorks Inc., USA) with a Gaussian function:

$f\left( x \right)=a1\times e^{\left( -\left( \frac{X-b1}{C1} \right)^{2} \right)}$

Where a1 is the maximum height of the Gaussian (I_max_), b1 – the peak position and c1 is the full width at half maximum (FWHM).

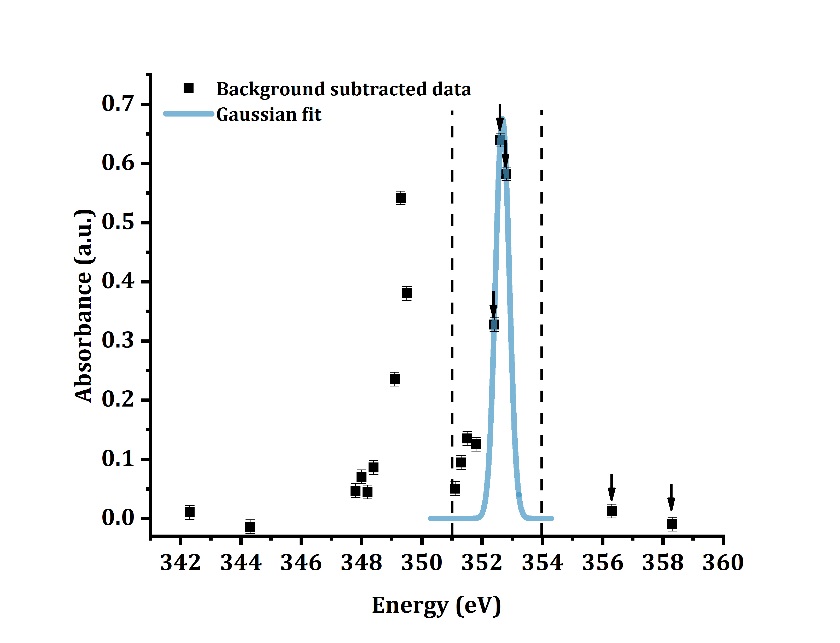

Fig S11. Gaussian fitting to the L_2_-peak data points. The background-subtracted data set is in black squares. The blue curve is the fitted Gaussian generated by Matlab. The Gaussian was fitted to the points marked by arrows.

The integrated absorbance (A) was calculated using the Gaussian fit results:

$$A={\sqrt{\pi}\times I}_{max}\times FWHM$$

Both I_max_ and the FWHM are taken as the average of the 95% confidence interval for each variable.
The concentration was calculated using:

$${\#Ca}_{{atoms/ Å}^{3}} =\frac{A}{\sigma\times0.67D}$$

$$\left[ Ca \right]\left( M \right)=\#Ca\frac{atoms}{Å^{3}}\times\frac{1Å^{3}}{{{(10}^{-10})}^{3}m^{3}}\times\frac{1m^{3}}{1000L}\times\frac{1mol}{6.022\times{10}^{23}atoms}$$

The uncertainties in I_max_ and FWHM values were taken as the half range of the 95% confidence intervals:

$$\Delta I_{max}={(I_{max})}_{\max95\% confidence}-I_{max,averaged}$$

ΔFWHM was calculated in a similar manner.
The fractional error of the integrated absorbance is then:

$$\frac{\Delta A}{A}=\sqrt{\left( \frac{\Delta I_{max}}{I_{max}} \right)^{2}+\left( \frac{\Delta FWHM}{FWHM} \right)^{2}}$$

The final uncertainty reported in the figure is thus:

$$\boldsymbol{\Delta}\left[ \boldsymbol{Ca} \right]\boldsymbol{=}\frac{\boldsymbol{\Delta A}}{\boldsymbol{A}}\boldsymbol{\times[Ca]}$$
